# An Anti-diabetic Nutraceutical enhances muscle cell glucose uptake of cardiometabolic drugs

**DOI:** 10.1101/2023.03.04.531136

**Authors:** K Saranya, RN Arpitha Reddy, Sanjana Battula, Ishita Mehta, Gopi Kadiyala, Subramanian Iyer, Subrahmanyam Vangala, Satish Chandran, Uday Saxena

**Author notes:** Address correspondence to: Dr. Uday Saxena.

## Abstract

Type 2 diabetes is currently treated by multiple drugs that are often combined to achieve maximum blood glucose lowering in patients. Yet more than 50% of patients are unable to attain target glucose levels. There is clear need for agents which can be added to current drugs to help patients achieve their target blood glucose levels safely.

To this end we tested an anti-diabetic nutraceutical for its ability to enhance glucose uptake in muscle cells in combination with currently used cardiometabolic drugs. We show here that the nutraceutical is able to effectively improve glucose uptake of multiple drugs suggesting that it may help in enhancement of glucose lowering by the drugs in patients and achieve optimal glucose levels.

## Introduction

Type 2 Diabetes is a condition that impairs the body’s ability to deliver and utilize blood glucose. High glucose blood levels are responsible for the morbidities and lifestyle changes associated with diabetes. Type 2 diabetes is a lifelong disease with severe consequences if blood glucose is left uncontrolled. It may lead to irreversible kidney damage (nephropathy), nerve damage (neuropathy), damage to eyes (retinopathy), heart disease and stroke. It is considered the worlds largest chronic disease with continuous increase in number of patients (1,2).

These are several oral drugs on the market today for glucose control. Some of these are metformin (acts on liver), Glimepiride and semaglutide (act on pancreas), Gliptins (act on an enzyme in blood) and combinations of these. Despite all of these drugs, in almost half of the patients, reaching healthy levels of blood sugar is still not possible (3,4). If the dose of the same drugs is increased in the hope of reaching lower glucose levels there is a risk danger of hypoglycaemia (blood glucose may become too low and there is risk of coma and damage to brain) and increased side effects.

Why are all these drugs alone or in combination not able to reduce glucose further? The answer may lie in the fact 70% of glucose lowering occurs due to uptake of blood glucose by the muscle. Yet, none of the present drugs have any affect directly on the muscle. Shown below in Table 1 are some of the current drugs and their target organs. As can be seen none of the drugs target glucose metabolism directly in the muscle.

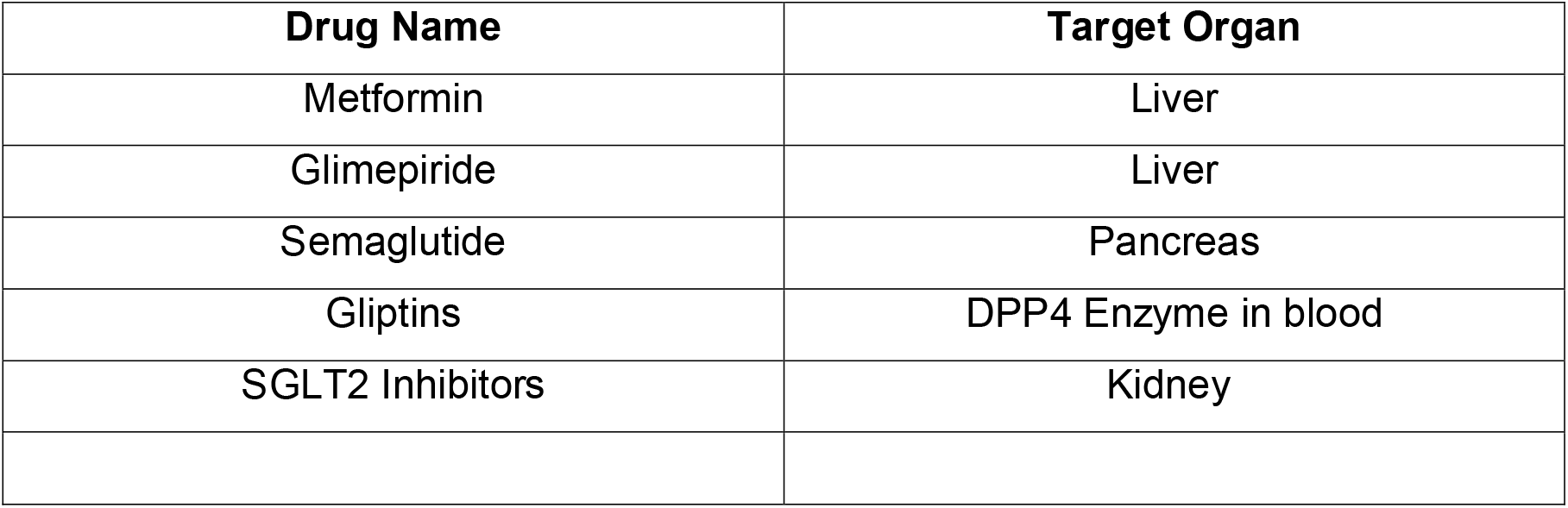

| Drug Name | Target Organ |
| --- | --- |
| Metformin | Liver |
| Glimepiride | Liver |
| Semaglutide | Pancreas |
| Gliptins | DPP4 Enzyme in blood |
| SGLT2 Inhibitors | Kidney |

To maximize glucose lowering we believe that any drug or nutraceutical has to impact muscle glucose metabolism. Insulin acts on the muscle via its receptor but increased dose of insulin has additional complications including hypoglycaemia. In view of this significant gap in current therapy with regards to targeting muscle biology, we designed a safe nutraceutical, DIVITIZ which can be added to current drug regimen and help achieve healthy blood glucose levels safely.

Our anti-diabetic nutraceutical specially targets molecular targets in the muscle glucose uptake and metabolism (5) and may provide additional glucose control over and above current anti-diabetic drugs.

We demonstrate here the nutraceutical can enhance glucose uptake by semaglutide and pioglitazone when added in combination in an in vitro muscle system. In addition the nutraceutical is also able to alleviate the known suppression of glucose uptake cause by statins (Lipitor/atorvastatin and simvastatin) thru a new mechanism of action. We propose that the nutraceutical DIVITIZ could be used in combination therapy with existing drugs to improve their glucose lowering efficacy.

## Methods

Pancreatic Min 6 cells, liver cell (Hepg2) and L6 Muscle cells, were used as and maintained in Dulbecco’s modified or Eagle’s Medium supplemented with 15% fetal bovine serum and 1% Penicillin/Streptomycin. The same media was also used for 3D bioprinting. A custom made 3D bioprinter with our own CAD program was used. Printing pressure for the cell layers and were optimized for each of the cell types in 96 well plates. Cells to be bioprinted were loaded in a 10 m l syringe Typically after printing each layer the plate was put back in incubator for 24-48 hrs and next layer was then printed.

To study the effects of a drug on glucose uptake, the drug was added to the 3D bioprinted layers and incubated at 37 C overnight. Next day, the cell layers are examined for cell morphology using a phase contrast microscope washed with media and then 10 uM of 2-(N-[7-nitrobenz-2-oxa-1, 3-diazol-4-yl] amino)-2-deoxyglucose (2-NBDG) (Molecular Probes-Invitrogen, CA, USA) was used to assess glucose uptake in the presence or absence of 100 nM insulin as the initiating step and incubated for 20 or 30 minutes. The data is reported as percent of control cells (no treatment, incubation with media).

Glucose uptake studies using L6 cells were performed similar to our previous reports (6). 2-NBDG glucose uptake was used to assess glucose uptake in L6 cells. Cells were kept in glucose free medium for half an hour before insulin stimulation. Cells were stimulated with 100 nM insulin for 20 and 30 min and incubated with 10 μ M of 2-NBDG for 15 min. The reaction was stopped by washing with cold PBS three times and the cells were lysed 0.1% Triton X. The lysate was then used to read florescence at 535 nM. The data is reported as percent of control cells (no treatment, incubation with media).

## Results

### 1. DIVITIZ improves insulin mediated uptake of glucose in 3D bioprinted diabetes model

We used a 3D bioprinted model of type 2 diabetes wherein all important cell types engaged in glucose metabolism were 3D bioprinted in the following sequence, liver cells on top followed by pancreas cells and finally the muscle cells. This system allows us to study drugs which may act on liver (metformin), pancreas (semaglutide) and muscle (pioglitazone). We first wanted to demonstrate that DIVITIZ and other drugs alone can enhance insulin stimulated uptake of glucose in this 3D system. As shown in Figure 1, metformin and semaglutide were not very effective in increasing glucose uptake relative to control (insulin stimulation but no drug added)). However pioglitazone and DIVITIZ were able to increase insulin stimulated uptake by 220% (30 minutes after insulin addition) and 150% (20 minutes after insulin stimulation) percent respectively. Therefore the 3D system is able to recapitulate pharmacological properties of some of the anti-diabetic drugs and DIVITIZ.

**Figure 1:**
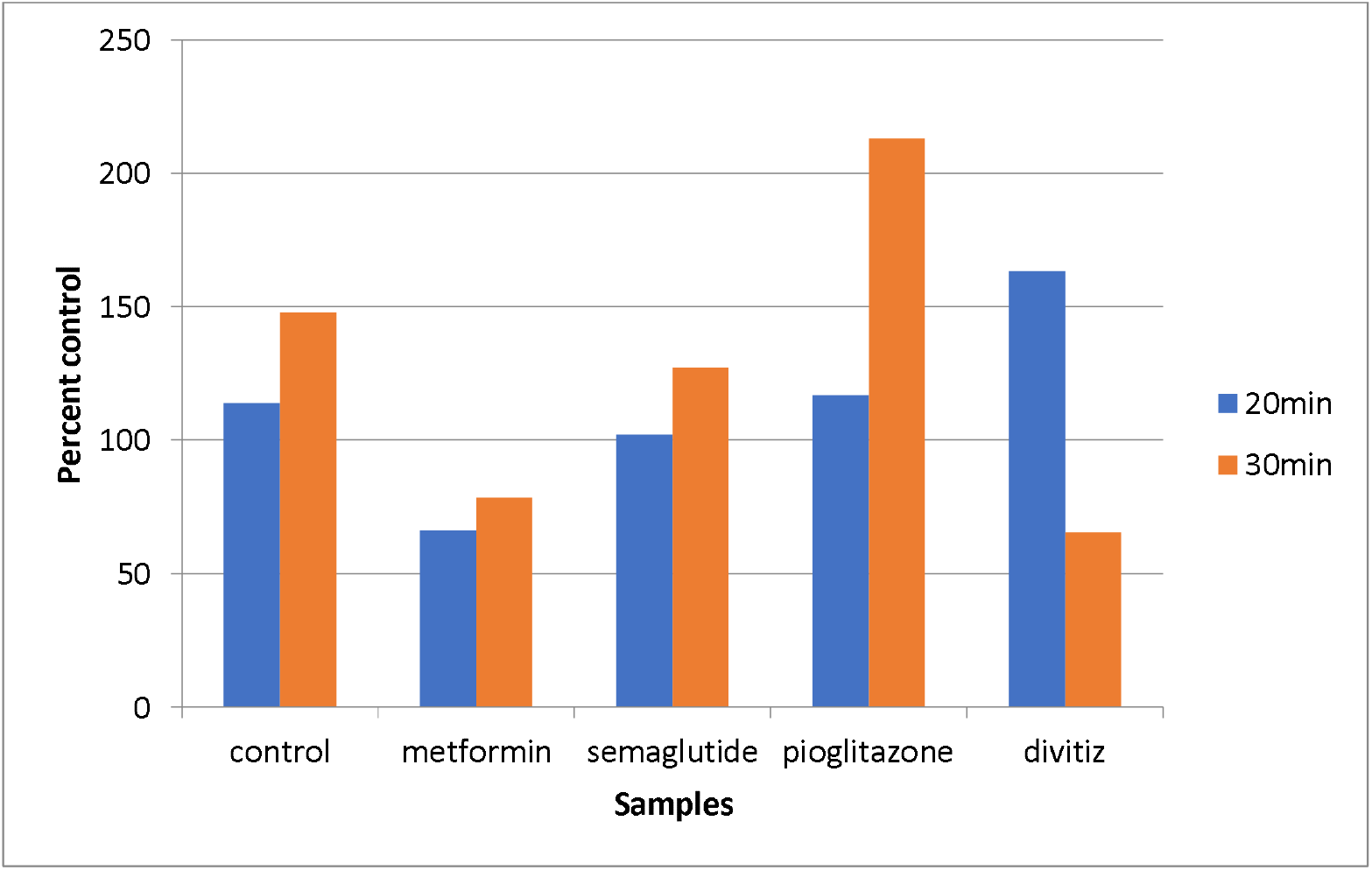
Effect of anti-diabetic drugs on glucose uptake in 3D model

### 2. DIVITIZ when added to anti-diabetic drugs further improves glucose uptake

We then explored if DIVITIZ would be able to enhance the uptake of above drugs when added in combination. Shown in Figure 2 is the data from those studies. The data show that DIVITIZ does not enhance glucose uptake by metformin but clearly enhances uptake in combination with semaglutide (over 200% at 20 minutes) and pioglitazone (about 150% at 30 minutes) relative to the drugs alone.

**Figure 2:**
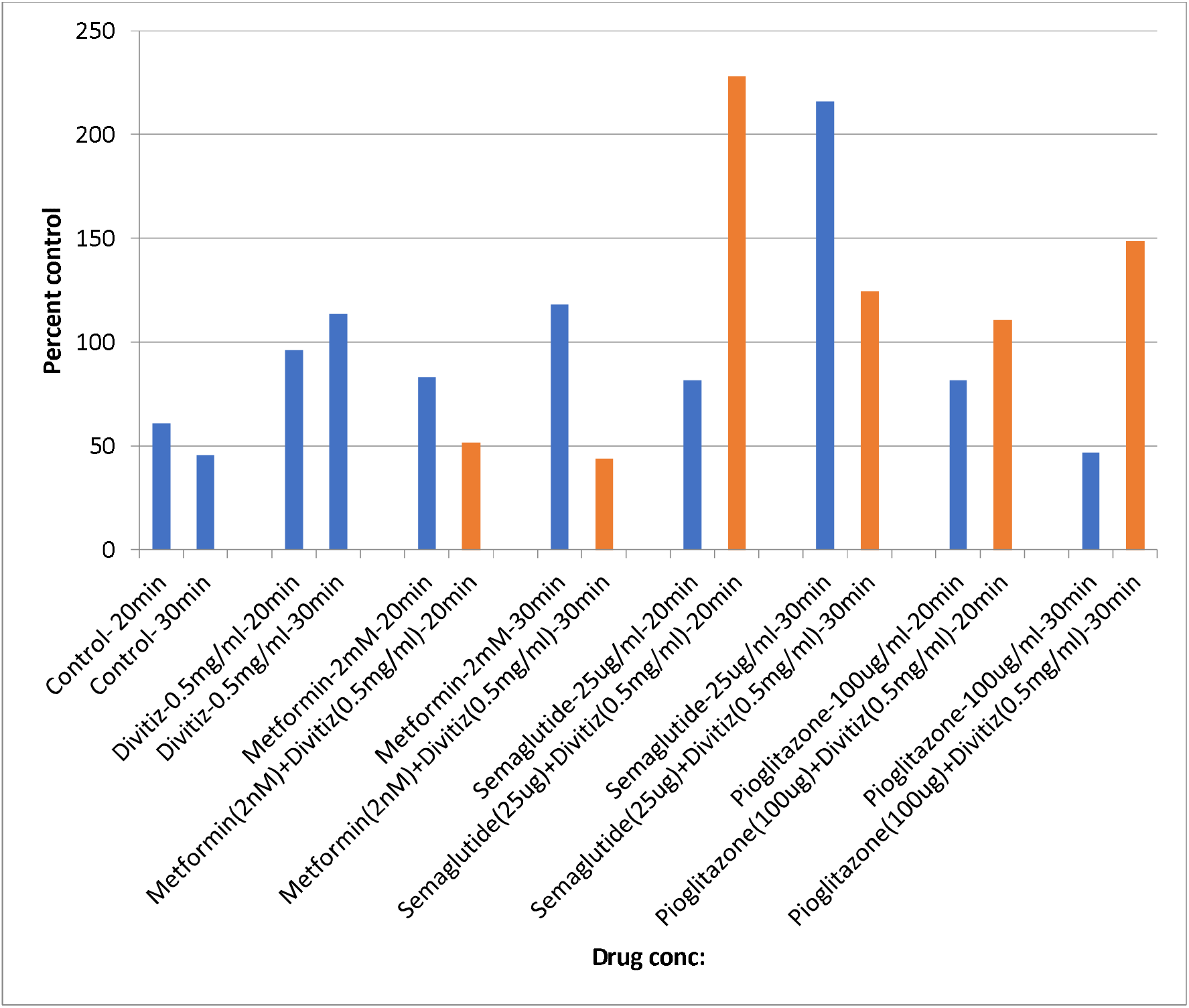
Effect of combining anti-diabetic drugs with DIVITIZ at various concentrations

Metformin works by inhibiting gluconeogenesis and therefore it is not surprising that we did not see an effect of combining it with DIVITIZ since the end point measured here in glucose uptake by muscle and not gluconeogenesis.

In combination studies with Semaglutide, we are able to demonstrate a massive increase with DIVITIZ. Similarly pioglitazone also showed an increase in uptake when combined with DIVITIZ. Collectively these data suggest that DIVITIZ has the potential to improve the efficacy of some of the anti-diabetic drugs.

### 3. DIVITIZ reverses statin induced suppression of glucose uptake

One of the well-established side effects of statins (cholesterol lowering drugs such as Lipitor and simvastatin) (6,7) is induction of insulin resistance which may ultimately lead to type 2 diabetes. We tested whether DIVITIZ could attenuate the inhibitory effects of lipitor and simvastatin on glucose uptake in an in vitro L6 muscle study. Shown in Figure 3 A and B are data showing that both lipitor and simvastatin reduced glucose uptake by 30 and 40% respectively. When the statins were added in combination with DIVITIZ, there was increased glucose uptake over statins alone suggesting that the nutraceutical is able to relive statin suppression. The reversal was dependent on concentration of DIVITIZ, especially at 0.25 mg/ml and 0.125 mg/ml added and the greatest reversal was shown at higher concentrations of DIVITIZ. The reversal by DIVITIZ was similar for lipitor and simvastatin.

**Figure 3 A:**
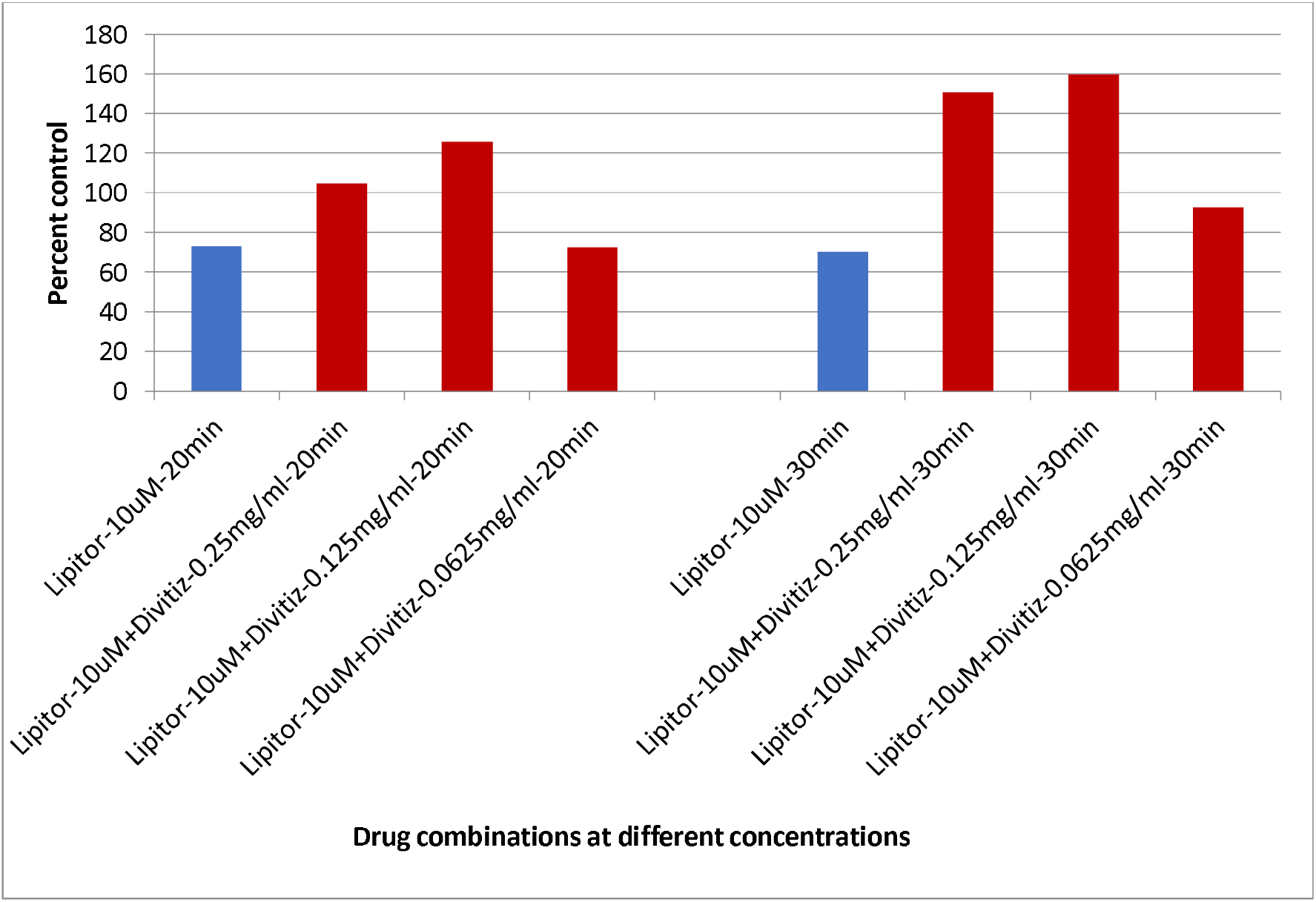
DIVITIZ alleviates suppression of glucose uptake by lipitor

**Figure 3 B:**
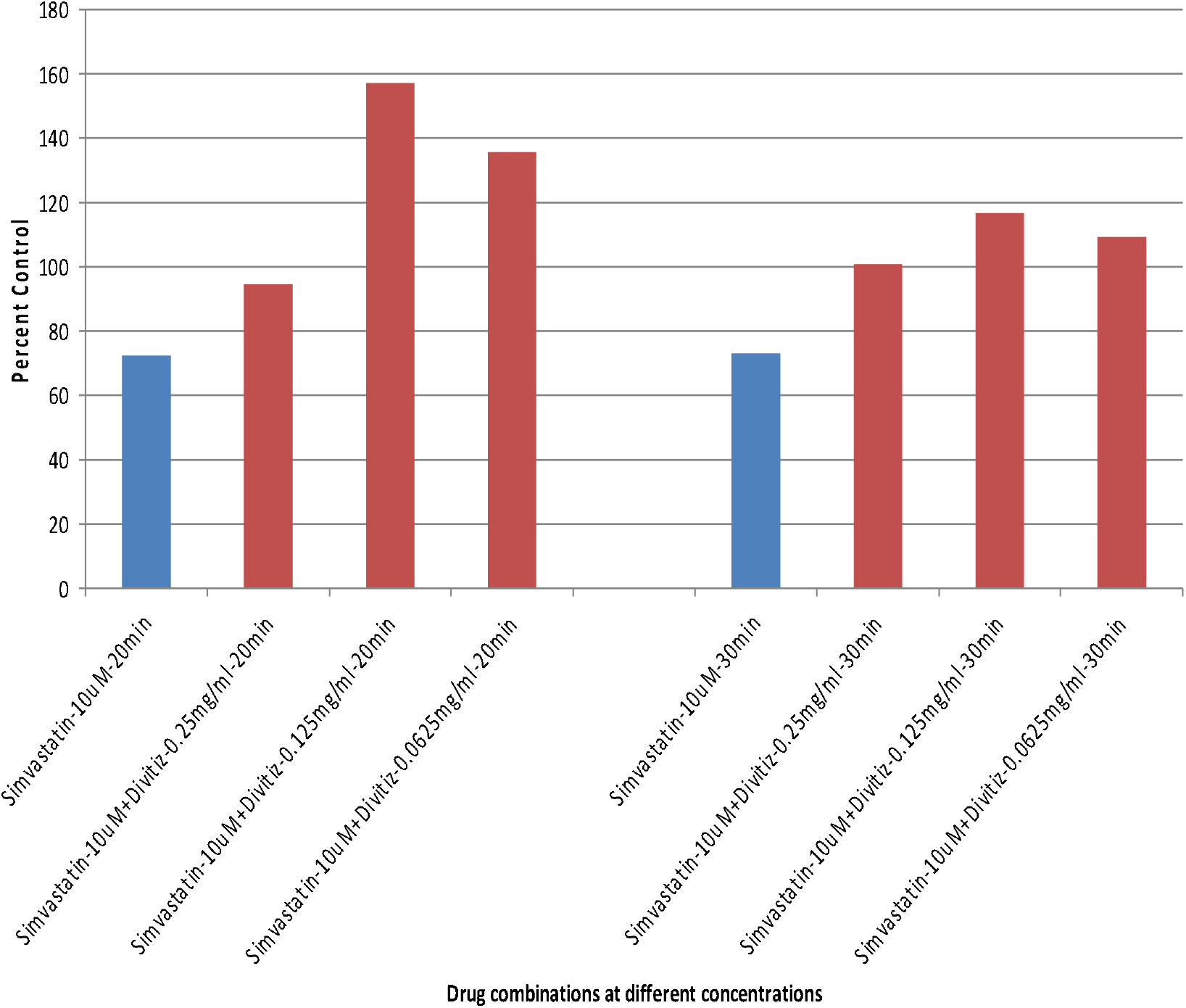
DIVITIZ relieves suppression of glucose uptake by simvastatin

### 4. DIVITIZ relieves statin suppression by new pathway

We then explored which of the three components of DIVITIZ individually may be important is relieving statin effects. We tested lipoic acid or niacinamide or Vitamin D3 in combination Lipitor in L6 cells. As shown below in figure 4 component niacinamide showed the biggest reversal at 20 and 30 minutes after insulin stimulation. Lipoic acid also showed reversal to a lesser extent. These data are suggestive that niacinamide depletion and redox stress are caused by Lipitor and therefore its suppression is reversed by the two components of DIVITIZ. It is important to reiterate here that previous reports have shown that Lipitor causes insulin resistance by accumulation of fatty acids in the cells (6). When the accumulated fatty acids are oxidized in the mitochondria, it may deplete niacinamide adenine dinucleotide (NAD) and caused redox stresses. As such when DIVITIZ components niacinamide and lipoic acid are added, the NAD depletion and redox stress are reversed and the glucose uptake suppression of Lipitor is overcome. This is the first demonstration of how Lipitor suppression of glucose uptake may be relieved pharmacologically.

**Figure 4:**
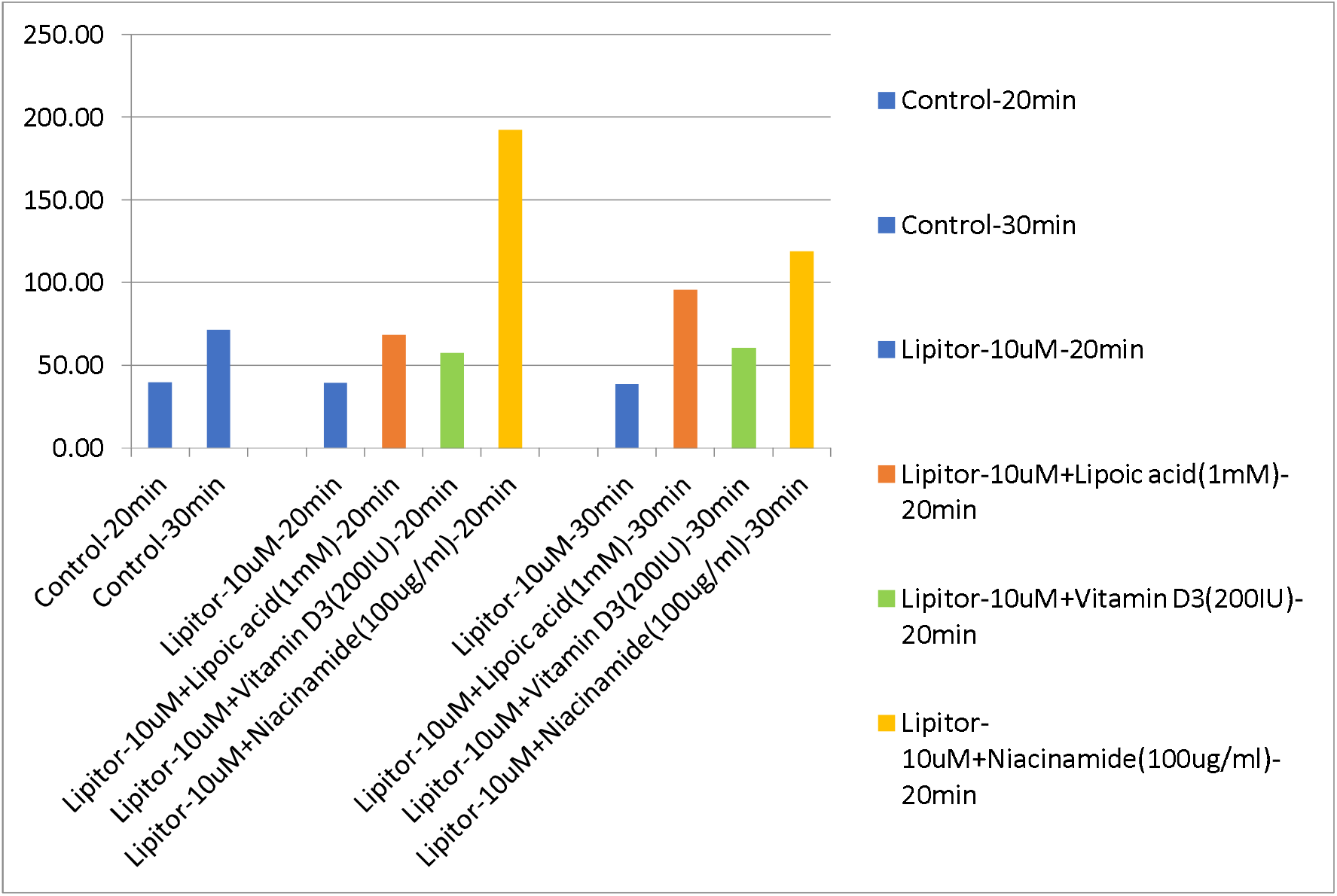
Effect of DIVITIZ components on Lipitor induced suppression of glucose uptake

## Discussion

### 1. Ability to enhance glucose uptake by anti-diabetic drugs and its consequences

Our data shown here demonstrate that the nutraceutical DIVITIZ is able to improve upon the glucose uptake caused by semaglutide and pioglitazone. The importance of this finding is that in clinical practice it may possible to achieve lower glucose levels by adding the nutraceutical to drug treatment regimen of anti-diabetic drugs. The option most easily available to physicians when target blood glucose levels are not met, is to increase the dose of existing drugs. However this may come with a price-metformin for example cannot be used at doses higher than 2 grams a day due to fear of lactic acidosis, semaglutide has the potential to cause thyroid C cell tumours and pioglitazone had a warning for pulmonary oedema. Thus if a safe nutraceutical such as DIVITIZ (composed of Vitamin D3, Niacinamide and lipoic acid in a fixed dose combination) can be used then it presents a whole new avenue for attaining target glucose levels without increasing doses of existing drugs. The results reported here if shown in clinical studies would be very significant for treatment of type 2 diabetes.

### 2. Ability to relieve statin suppression and its clinical implications

Statin family are lifesaving cholesterol lowering drugs that have been used globally for primary and secondary prevention of cardiovascular disease. As much as they are very successful drugs, recent evidence suggests that they impair muscle cells ability to utilize glucose and increase propensity for type 2 diabetes (6,7). Thus the physician has a conundrum, because on one hand they are used to treat cardiovascular disease and on the other hand they may induce type 2 diabetes which in itself is a risk factor to cardiovascular disease. Our finding of DIVITIZ relieving statin suppression of muscle glucose uptake therefore presents an opportunity for the patients i.e. statins could be taken together with DIVITIZ to minimize insulin resistance. Future clinical trials are needed to establish the benefits of this combination.

## Notes

### Competing Interest Statement

The authors have declared no competing interest.

